# Lateral Subunit Coupling Determines Intermediate Filament Mechanics

**DOI:** 10.1101/676197

**Authors:** Charlotta Lorenz, Johanna Forsting, Anna V. Schepers, Julia Kraxner, Susanne Bauch, Hannes Witt, Stefan Klumpp, Sarah Köster

## Abstract

The cytoskeleton is a composite network of three types of protein filaments, among which intermediate filaments (IFs) are the most extensible ones. Two very important IFs are keratin and vimentin, which have similar molecular architectures, but different mechanical behaviors. Here we compare the mechanical response of single keratin and vimentin filaments using optical tweezers. We show that the mechanics of vimentin strongly depends on the ionic strength of the buffer and that its force-strain curve suggests a high degree of cooperativity between subunits. Indeed, a computational model indicates that in contrast to keratin, vimentin is characterized by strong lateral subunit coupling of its charged monomers during unfolding of *α*-helices. We conclude that cells can tune their mechanics by differential use of keratin versus vimentin.

---

The cytoskeleton is composed of three types of biopolymers – actin filaments, microtubules and intermediate filaments (IFs) – which, along with cross-linkers and motor proteins, form a dense network in the cell [1], and determine its mechanical properties. Microtubules and actin filaments are conserved across different cell types and organisms. By contrast, IFs are expressed in a cell-type specific manner [2–4]: Thus, for example, keratins are predominantly expressed in epithelial cells and vimentin in cells of mesenchymal origin. It has been shown that vimentin deprived cells are less mechanically stable and migrate more slowly [5], whereas cells lacking keratin are softer and more deformable [6, 7]. These differences are likely to play an important role during endothelial-to-mesenchymal transition (EMT), for example in embryogenesis, wound healing and cancer metastasis, when cells upregulate vimentin expression and downregulate keratin expression [4, 8–10]. We hypothesize that keratin and vimentin IFs have different mechanical properties already at the single filament level. It has been shown previously that single vimentin IFs exhibit a pronounced extensibility of up to 4.5 times their original length [11–13] and a high flexibility [14, 15], and can dissipate up to 80% of the input energy when stretched and relaxed [16]. Keratin has so far been primarily studied in the context of bundles, for example in hagfish slime threads [17], wool fibers [18] and hard *α*-keratin fibers [19]. However, data from single filaments are needed to decouple the mechanics resulting from the bundle or network structure, i.e. the inter-filament interactions, from the single filament mechanics.

IF mechanics are closely linked to the their molecular architecture [13, 16, 20]. The monomer consists of a “rod” domain including three *α*-helices, which are connected by linkers and flanked by intrinsically disordered head and tail regions (Fig. 1a,b) [21]. Despite differing amino acid sequence, all cytoskeletal IFs share this monomer structure, as well as the particular assembly pathway: Two monomers form a parallel dimer, two dimers an antiparallel, half-staggered tetramer and tetramers eventually form unit-length-filaments (ULFs) with a length of about 60 nm [21]. This lateral assembly is followed by longitudinal annealing of ULFs resulting in *µ*m-long filaments. One important difference between keratin and vimentin IFs is the average number of tetramers per filament cross-section of four and eight, respectively [2]. During stretching of these filaments, the *α*-helices open into a *β*-state leading to a contour length change [16, 22].

**FIG. 1.**
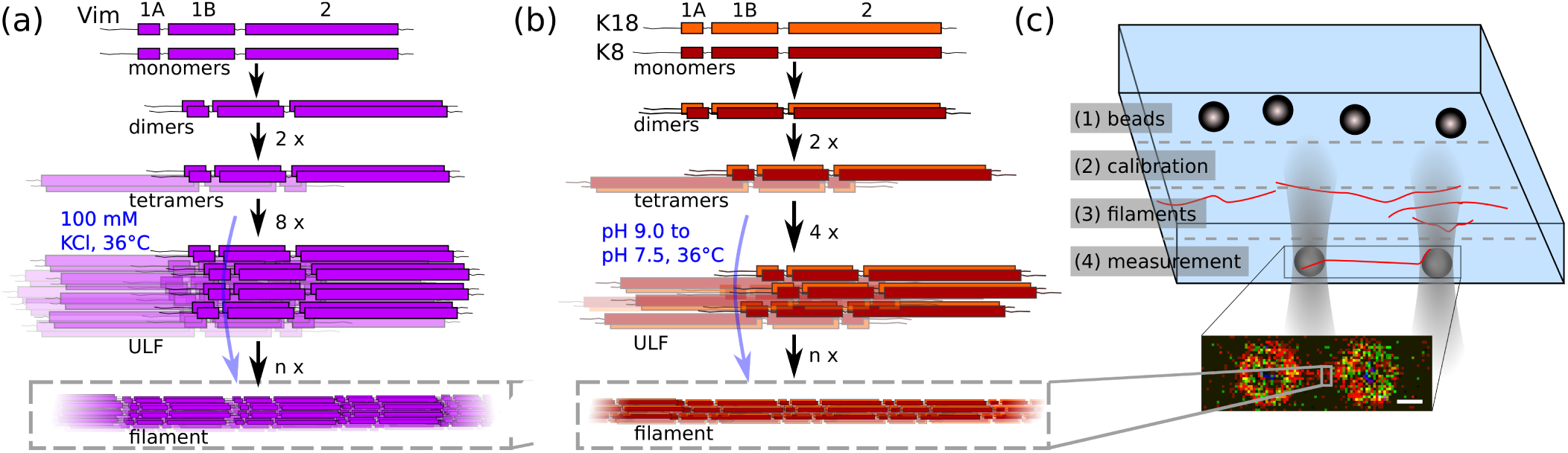
(a, b) Assembly pathway of vimentin (Vim) and keratin (K8, K18) IFs, respectively. The monomers consist of three *α*-helical regions (1A, 1B, 2) connected by two linkers and flanked by intrinsically unstructured regions, and form extended IFs in a strictly hierarchical manner. (c) Top: Schematic of the microfluidic device used for measurements with the OT. Bottom: Confocal image of a fluorescently labeled keratin IF captured between 4.42 *µ*m-diameter-beads; scale bar 2 *µ*m.

Here, we study the mechanical behavior of single keratin and vimentin IFs under load by stretching them with optical traps (OTs) [13, 16]. It is well known that keratin and vimentin are held together mainly by hydrophobic and electrostatic interactions. Therefore, we use two distinctly different buffer conditions with high or low ionic strength, respectively, to tune the electrostatic interactions. We find that ionic strength impacts IF mechanics, which we explain by stronger lateral coupling in vimentin subunits than in keratin subunits, corroborated by data from atomic force microscopy (AFM). The experimental data from OT are modelled and quantitatively fitted by a Monte Carlo (MC) simulation based on the IF structure [16]. From the fit, we obtain the free energy difference between the folded *α*-helix and the unfolded state, and the *α*-helical stiffness.

Human keratin 18 (K18), keratin 8 (K8), K8 with an additional cysteine at the C-terminus (Cys-K8) and the vimentin mutant C328A with an additional cysteine at the C-terminus (Cys-Vim) were recombinantly expressed, labeled and reconstituted to tetrameric form as described in the SI (URL inserted by publisher). Keratin (50% K18, 25% K8, 20% Cys-K8, 5% labeled Cys-K8) was assembled at 0.1 g/L by dialysis into low ionic strength buffer (LB: 10 mM TRIS, pH 7.5) [23] and vimentin at a protein concentration of 0.2 g/L into high ionic strength buffer (HB: 100 mM KCl, 2 mM PB, pH 7.5) [15], both at 36 °C over night. In both cases about 4% of all monomers were labeled fluorescently with ATTO647N.

For OT measurements, the assembled keratin and vimentin filaments were diluted 1:70 and 1:100, respectively, with the corresponding assembly buffer (HB or LB). The OT trap setup (LUMICKS, Amsterdam, Netherlands) was equipped with a confocal fluorescence microscope and a microfluidic device as sketched in Fig. 1c. Polystyrene beads (Kisker Biotech, Steinfurt, Germany) were maleimide coated [24] to allow for covalent binding to the IFs *via* the additional cysteine in the monomers. In total, our experiment required four microfluidic subchannels: a bead channel (1), a buffer channel for calibration (2), an IF channel (3) and a channel for measurements (4). The beads were diluted with the assembly buffer of the respective studied protein (HB or LB), which was also used as buffer in the calibration channel. The buffer in channel (4) was either LB or HB and each IF was studied in both buffers.

Before each measurement, two beads were captured with the OT in channel (1) and the trap stiffness was calibrated *via* their thermal noise spectrum in channel (2). IFs were attached to the beads in channel (3) and it was ensured by fluorescence microscopy that only one IF bound to the beads. The traps with the IF were moved to channel (4) and incubated for 30 s, unless the measurement was intended to take place in the assembly buffer of the respective IF protein. One OT was moved with speeds between 0.3 *µ*m/s and 2.5 *µ*m/s to stretch the IF in channel (4). The force exerted on the IF by the OT as well as the bead positions were recorded.

IF heights were measured with a commercial AFM (Infinity, Oxford Instruments Asylum Research, Santa Barbara, CA). IFs were incubated for 30 s in the buffer of interest, fixed with 0.125% glutaraldehyde and imaged on a piece of silicon wafer (Crystec, Berlin, Germany) in buffer. Cantilevers (MLCT, Bruker, Billerica, MA, USA) were calibrated *via* their thermal noise spectrum.

From the OT data, the strain *ε* = *L/L*_0_−1 is calculated [16] using the measured IF length *L* and the IF length *L*_0_ at 5 pN. The individual (thin lines) and average (thick lines) force-strain curves of single keratin and vimentin IFs in the two buffer conditions are shown in Fig. 2a,b. The averages are calculated by averaging both force and strain data; this procedure is explained in detail in the SI (URL inserted by publisher).

**FIG. 2.**
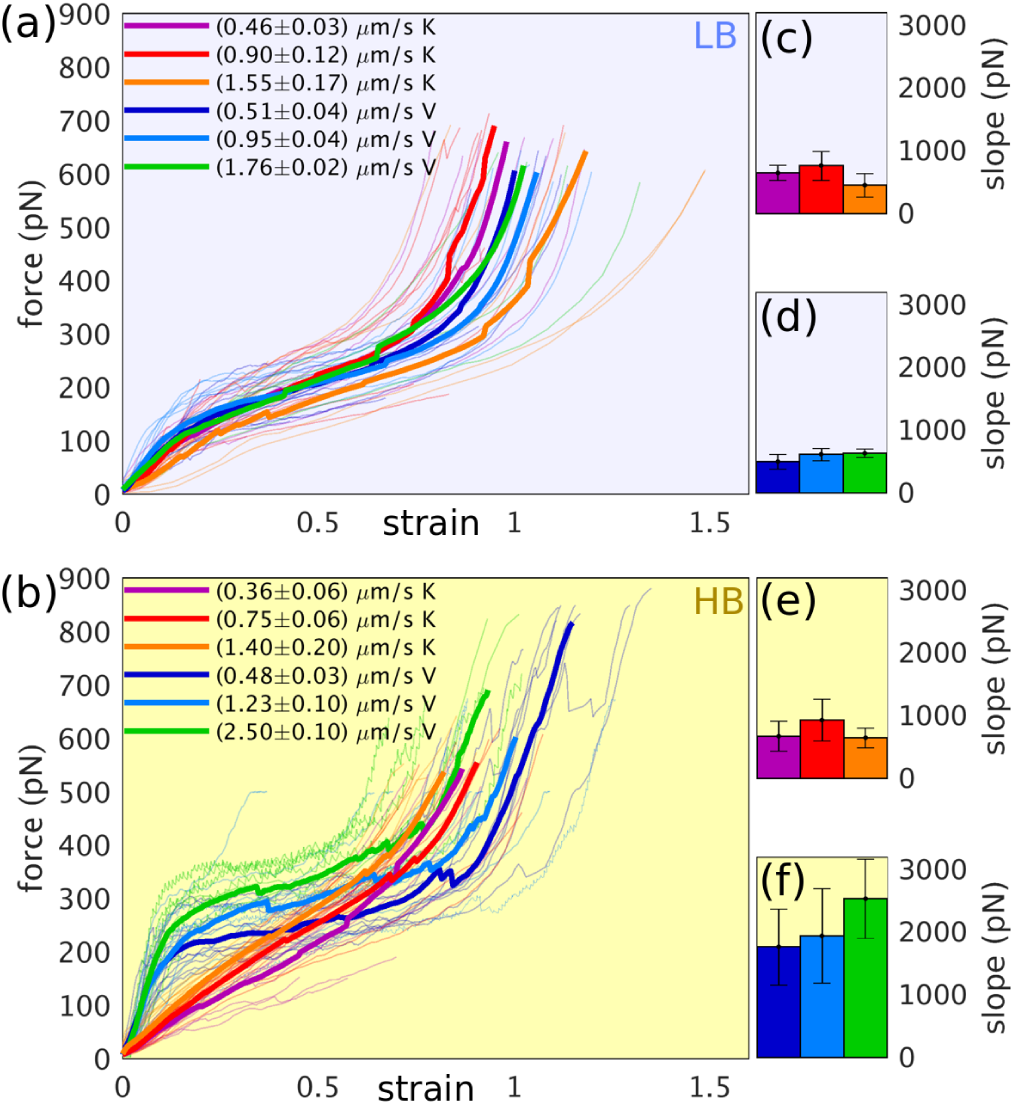
(a,b) Force-strain curves for keratin (K, warm colors) and vimentin (V, cold colors) in (a) LB (blue background) and (b) HB (yellow background) measured with the OT. The curves from single IFs (thin lines) for different loading rates (see color code) are averaged (thick lines). (c-e) Slopes from linear fits in the low strain regime of (c) keratin in LB, (d) vimentin in LB, (e) keratin in HB and (f) vimentin in HB.

In contrast to keratin, the force-strain behavior of vimentin filaments significantly depends on the ionic strength of the buffer as Fig. 2 shows. In LB, the force-strain behavior of keratin and vimentin is similar and can be divided into three regimes [13, 16]: There is an elastic regime for low strains caused by the elastic behavior of *α*-helices [13, 20, 25, 26]. A less steep regime for strains between 0.2 and 0.8 arises from the step-wise opening of *α*-helices during elongation [20, 22]. The filaments stiffen again for high strains since most *α*-helices are unfolded and the resulting structure is stretched [20]. The slopes for low strains (in the range between 0.015- or 0.015-0.15, depending on the linear regime) are on average slightly higher for vimentin compared to keratin filaments (Fig. 2c,d), which can be partially explained by the doubled number of monomers per cross-section in vimentin IFs. In summary, for keratin and vimentin filaments, there is no clear plateau-like regime in LB, similar to the stress-strain behavior of hagfish slime threads [17] (Fig. 2b). By contrast, both filament types behave differently in HB. For keratin filaments, there is no clear separation between the regimes. Vimentin filaments, however, show a distinct plateau-like force-strain behavior, which indicates some degree of cooperativity. Both proteins develop a loading-rate depending behavior and are stiffer in HB (Fig. 2e,f).

The different curve shapes for keratin and vimentin filaments also result in a higher input energy *E* for vimentin than for keratin filaments in HB. *E*_XY_ is calculated by integrating the force-strain curves up to a force of 500 pN of protein X in buffer Y. In LB, *E* of both protein filaments is the same, *i.e.* the ratio of the input energies *E*_VL_*/E*_KL_ is 1.01 ± 0.16, whereas in HB, vimentin filaments take up about 45% more energy (*E*_VH_*/E*_KH_ = 1.45 ± 0.25), as shown in the SI (URL inserted by publisher).

To understand these data, we take a closer look at the molecular properties of keratin and vimentin as we observe that the different ionic strength of the two buffers has a strong impact on the substructure of the filaments. Keratin monomers are 1.5 times more hydrophobic than vimentin monomers [27], so that the hydrophobic effect is stronger for keratin subunits than for vimentin, independent of electrostatic effects. Additionally, vimentin monomers are more negatively charged than keratin (19 *e*/monomer *vs.* 8.5 *e*/monomer for keratin). Concerning the buffers, HB has an about 20 times higher ionic strength than LB, which allows for a closer arrangement of the subunits in the filament since the ions screen the negative IF charges and lower the electrostatic repulsion of the subunits within the filament [28, 29]. These additional attractions need to be overcome when the filament is stretched, so that the filaments appear stiffer in HB. The lateral coupling between the keratin subunits becomes slightly stronger in HB than in LB, which reduces the maximum strain, raises the initial slope and renders the curve rather linear. Keratin monomers are less negatively charged so that the subunits do not repel each other as strongly as in vimentin. Therefore, we can attribute the larger stiffening and the plateau-like regime of vimentin filaments to a stronger attraction and a higher lateral coupling strength between the subunits. The plateau-like behavior also leads to a larger input energy as reported above. Consequently, vimentin IFs are able to absorb more energy under load than keratin IFs.

To test our hypothesis of the difference in lateral coupling strength of subunit attractions, we measure the height of keratin IFs and vimentin IFs each in both LB and HB by AFM. The attraction between the substrate and the IF flattens the originally circular cross-section of the filament [11, 15]. However, a stronger attraction between the filament subunits to each other prevents this effect. IFs in the AFM images are tracked and the IF height is extracted from the data as described in the SI (URL inserted by publisher). A typical AFM image is shown in Fig. 3a and the filament heights agree with literature [2, 30]. The average height for keratin IFs increases from LB to HB by a factor of 1.2 ± 0.4 and the height of vimentin IFs by a factor of 2.6 ± 0.9 (Fig. 3b,c). This supports our hypothesis that HB enhances the attractions between single subunits more strongly in vimentin filaments.

**FIG. 3.**
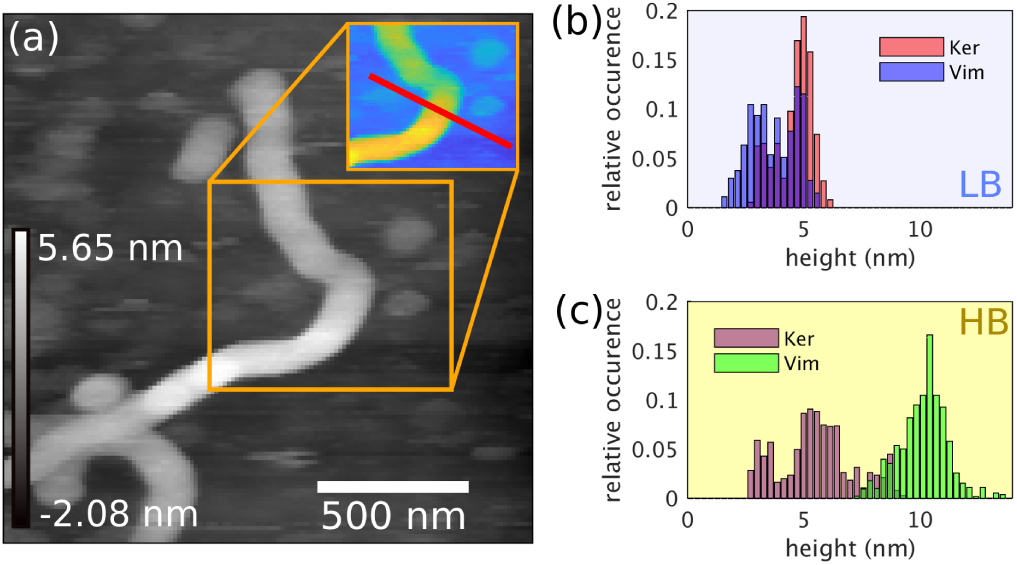
(a) Typical AFM image; the inset shows the processed AFM image in MatLab used to extract the height profile (red line). (b,c) Histogram of keratin and vimentin filament heights measured with AFM in (b) LB and (c) HB.

To understand why the lateral coupling in keratin and vimentin IFs has a different effect on the mechanical properties, *i.e.* why vimentin IFs exhibit a clear plateau in comparison to keratin IFs, we model keratin and vimentin IFs by a mechanic model which is based on Refs. [16, 31, 32]: Each monomer is described as a spring in series with an element that can elongate under tension (Fig. 4a,b). The spring corresponds to the elastic behavior of an *α*-helix for low forces. The energy difference between the *α*- and *β*-state is Δ*G*. Before stretching, *N*_*P*_ monomers are connected in parallel as an equivalent to a ULF (*N*_*P*_ = 16 for keratin, *N*_*P*_ = 32 for vimentin). To model a filament, 100 of these ULFs are connected in series by springs which sum up all linkers between the ULFs. The force-strain behavior of the modeled filament is determined by a MC simulation. With respect to the elongation of the filament, two variants of the model are simulated: In the first (uncoupled) case (1), the filament elongates, when all monomers in one subunit are in the *β*-state (Fig. 4a), in the second (coupled) case (2), the filament elongates, when all monomers of one ULF are in the *β*-state (Fig. 4b) [16]. Thus, case (1) supports the idea of protofilaments within the filament which can slide past each other [33–35].

**FIG. 4.**
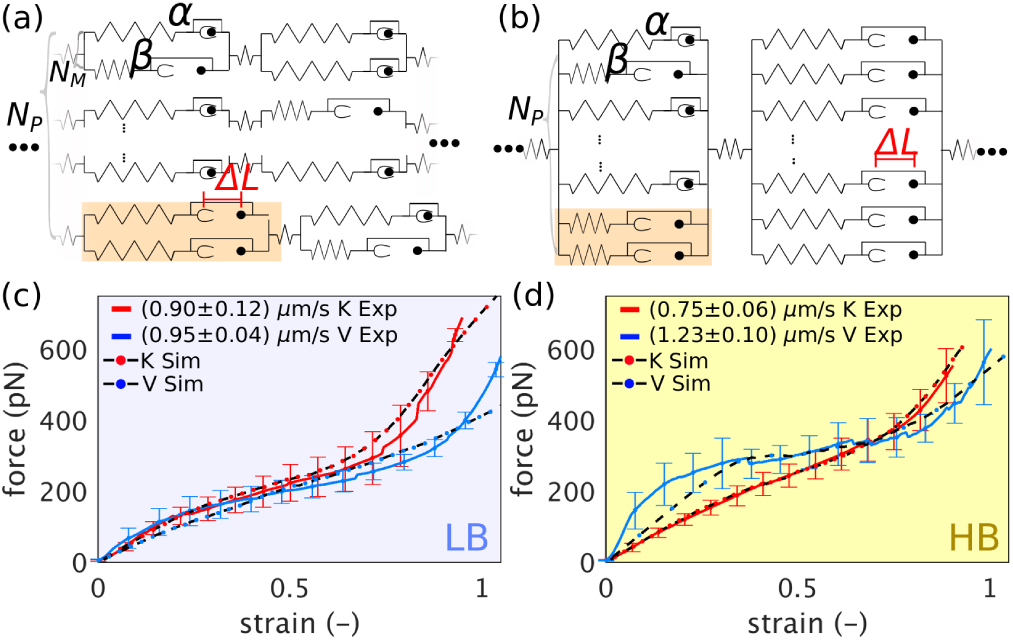
(a) Model for uncoupled dimers as subunits (case 1) and (b) model for coupled dimers as subunits (case 2). If all subunits are in the *β*-state in case 1, this leads to an elongation by Δ*L*, whereas in case 2 all monomers have to be in the *β*-state for elongation (orange: elements under discussion in the text). (c,d) Simulated and measured force-strain curves for keratin (K) and vimentin (V) in (c) LB and (d) HB.

The coupled and uncoupled extensions differ in how the force *ϕ* is shared among the monomers *M* and thus in the force dependence of the transition rates 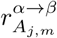 from the *α*- to the *β*-state (for details see the SI (URL inserted by publisher)). The transition rate has the general form:

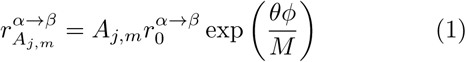

with the number of monomers *A*_*j,m*_ in the *α*-state of the *m*th subunit in the *j*th ULF, the zero-force reaction rate from a monomer in the *α*- to the *β*-state 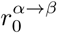 and the load distribution factor *θ* [36, 37]. In the uncoupled case, the force is shared equally among the subunits and within a subunit among the monomers, *M* = *N*_*C*_ *A*_*j,m*_, while in the coupled case, the force is shared equally among all monomers of the ULF,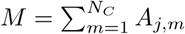 (with the number of laterally associated subunits *N*_*C*_).

The two different assumptions for elongation lead to a fundamentally different force-strain behavior: In case (1), the data are smooth and “s-shaped”, since the laterally associated *α*-helices in the subunits can open independently at a certain minimum force. In case (2), a plateau evolves, because a higher minimum force is needed to open the laterally coupled *α*-helices. Once that force is reached, the *α*-helices in one ULF open in a cascading manner, which is also observed by Erdmann and Schwarz in Ref. [32] for a system that resembles one vimentin ULF in HB in our model. We observe a plateau only for vimentin filaments in HB, thus, we model them with case (2) so that all *α*-helices in the ULF have to be unfolded for a length extension [13, 16]. For keratin filaments in HB and vimentin and keratin filaments in LB, we assume case (1) which corresponds to an uncoupled filament elongation. A self-written MatLab code fits the simulation data to the experimental data obtained with OT (Fig. 4). The simulations agree well with the experimental force-strain curves for keratin filaments in LB and HB and for vimentin filaments in HB. The experimental data of vimentin filaments in HB exhibit a higher initial slope than expected from the simulation. In both buffers, Δ*G*, as extracted from the fit parameters lies around 0.3 or 0.6 *k*_*B*_*T* per amino acid for keratin or vimentin, respectively, and agrees with theoretical results for comparatively short peptides (2-37 amino acids) from literature [38–40]. This indicates that the energy stored in a single *α*-helix does not depend on the ion concentration, in contrast to the lateral coupling strength between the *α*-helices. We also determine the *α*-helical stiffness of about 0.6-3.4 pN/nm from the simulation parameters in agreement with literature [13, 41, 42].

Our data from OT, AFM and MC simulations strongly indicate that the lateral coupling in vimentin filaments induced by additional cations is so strong that all parallel *α*-helices in one ULF have to unfold to induce a length change, whereas in keratin filaments the filament elongates in any condition as soon as one subunit is in the unfolded configuration. Hence, vimentin filaments are able to absorb more mechanical energy than keratin. We assume that three main differences in the physical and molecular properties of the two IFs contribute to the different lateral coupling strengths: (i) electrostatics, (ii) hydrophobicity and (iii) compaction. Aspects (i) and (ii) have been discussed above. Concerning aspect (iii), in contrast to keratin, vimentin IFs compact after elongation [43, 44]. Data from hydrogen-deuterium exchange show that for this compaction, charged amino acids in linker L1 and in coil 1B attract oppositely charged amino acids in linker L12 from a neighboring tetramer, which causes an additional link between these two tetramers [44]. Therefore, compaction can additionally increase the lateral coupling strength in vimentin IFs. Keratin IFs do not compact [45] and also do not exhibit the same charge pattern that all compacting IFs have in common in the concerned sequence [46] (see SI (URL inserted by publisher) for the detailed amino acid overview). Note that it is not the different number of monomers per cross-section in keratin and vimentin that causes the completely different behavior in HB, as an uncoupled filament with 16 monomers in the MC simulation does not exhibit a plateau either (see the SI (URL inserted by publisher) for a comparison).

To conclude, our experiments substantiate the idea that cells can fine-tune their ability to absorb large amounts of energy to protect the cell from mechanical damage by the expression of different IFs. Remarkably, our measurements on the *µ*m-scale allow us to conclude upon interactions of the filament subunits on the nm-scale. Depending on the surrounding ion concentration, vimentin filaments stiffen and absorb more energy than keratin filaments. An MC simulation based on assumptions about the molecular structure of both IFs show that a stronger lateral coupling of the vimentin subunits leads to a plateau-like regime and a higher absorbed energy.

We thank I. Mey for support with the AFM measurements and data analysis. We are grateful for fruitful discussions and technical support by H. Herrmann, A. Janshoff, U.S. Schwarz, J. Kayser, N. Mücke and P. Rauch. The work was financially supported by the European Research Council (ERC, Grant number CoG 724932) and the Studienstiftung des deutschen Volkes e.V.

