## Supplementary Material for "Lateral Subunit Coupling Determines Intermediate Filament Mechanics"

### Supplementary Information: Lateral Subunit Coupling Determines Intermediate Filament Mechanics

Charlotta Lorenz[1], Johanna Forsting[1], Anna V. Schepers[1], Julia Kraxner[1], Susanne Bauch[1], Hannes Witt[2,3], Stefan Klumpp[4], and Sarah Köster[1*]
[1]*Institute for X-Ray Physics, University of Goettingen, Friedrich-Hund-Platz 1, 37077 Göttingen, Germany*
[2]*Institute for Organic and Biomolecular Chemistry, University of Goettingen, Tammanstraße 2, 37077 Göttingen, Germany*
[3]*Max-Planck-Institut für Dynamik und Selbstorganisation, Am Faßberg 7, 37077 Göttingen and*
[4] *Institute for Dynamics of Complex Systems, University of Goettingen, Friedrich-Hund-Platz 1, 37077 Göttingen, Germany*

#### MATERIALS AND METHODS

##### Protein expression, labeling and dialysis:

**Keratin:** Human keratin 18 (K18), keratin 8 (K8) and K8 with an additional cysteine at the C-terminus (Cys-K8) were recombinantly expressed in E. Coli [1]. The Cys-K8 was labeled with ATTO647N (AttoTech GmbH, Siegen, Germany) *via* maleimide bonding [2]: 1 mL of Cys-K8 at 1 mg/mL was dialyzed (Spectra/Por 25 kDa, Spectrum Laboratories, Piscataway, NJ, USA) into 5 M urea, 50 mM phosphate buffer, pH 7.0 (labeling buffer) over night. 20 $\mu$L of 10 mM ATTO647N, dissolved in DMSO, were added to the dialyzed protein in 5 $\mu$L portions with an incubation time of 5 min each. 100 $\mu$L of 1 M L-cystein (Carl Roth, Karlsruhe, Germany) were added to bind free dye molecules. After 1 h of incubation, the labeled protein was separated from the free dye by size exclusion chromatography (Bio-Gel P-30, Bio-Rad, München, Germany) with a 27 mL column [2]. The labeled protein was washed through the column by adding labeling buffer. The protein concentration was measured by UV/Vis-spectroscopy (NanoDrop One/OneC, ThermoFisher, Schwerte, Germany). Afterwards, the labeled Cys-K8 was dialyzed to 8 M urea, 50 mM TRIS, pH 9.0, (storage buffer) [2] and stored at -80 °C.

For preparation of assembly, K18 (50%), K8 (25%), unlabeled Cys-K8 (20%) and labeled Cys-K8 (5%) were dialyzed at a total protein concentration of 0.1 g/L in a stepwise manner (8 M, 6 M, 4 M, 2 M, 1 M urea) from storage buffer to 2 mM TRIS, pH 9.0, over night [3]. Keratin filaments were assembled by dialyzing the protein mixture at 0.1 g/L to 10 mM TRIS, pH 7.5, (LB) at 36 °C over night [3]. About 4% of all monomers had an additional ATTO647N molecule attached to the C-terminus.

**Vimentin:** Vimentin C328N with two additional glycines and one additional cysteine at the C-terminus was recombinantly expressed and labeled with ATTO647N as in Ref. [4]. Vimentin monomers were dialyzed in a stepwise manner (8 M, 6 M, 4 M, 2 M, 1 M urea) from 8 M urea, 50 mM phosphate buffer (PB), pH 7.5, to 2 mM sodium PB, pH 7.5, and assembled by dialysis at 36 °C over night to 100 mM KCl, 2 mM PB, pH 7.5 (HB) [5]. About 4% of all vimentin monomers were labeled with ATTO647N.

**AFM sample preparation:** The assembled protein was diluted 1:10 in the case of vimentin and 1:5 in the case of keratin with LB or HB and incubated for 30 s. Glutaraldehyde (2.5% in PBS) was prepared at a final concentration of 0.25% by dilution with the buffer that the protein was studied in (LB or HB). The diluted glutaraldehyde and the diluted protein were mixed 1:1 and incubated for 30 s. The mixture was transferred to a piece of silicon wafer (Crystec, Berlin, Germany) 0.8 cm × 1.2 cm and incubated for 3 to 5 min. 100 $\mu$L of fresh buffer were added three times and removed again to avoid non-adhered filaments on the silicon wafer. About 150 $\mu$L of buffer were left on the sample during imaging. The smallest tip on the multi-tip cantilever (MLCT, Bruker, Billerica, MA, USA) with a tip radius of about 20 nm was used for imaging in tapping mode .

#### DATA ANALYSIS

All data analysis and image processing is carried out with self-written MatLab codes.

**Averaging of force-strain curves:** Average force-strain curves are calculated by averaging the force and strain data for all curves in three steps as shown for typical single data sets in Fig. S1: (1) The force-strain data sets are classified as stable if the measured force exceeds 500 pN. For the curves in Fig. S1a, one filament is not stable and two are stable. (2) The force data from the stable filaments are interpolated to the same number of data points (200). The average of the maximum strain values of all stable filaments is calculated and set to the maximum strain for all stable filaments as shown in Fig. S1b. The strain data consist of 200 equally spaced data points from 0 up to the average maximum strain. The force data of all stable filaments are averaged for each strain value to obtain the average force-strain curve of all stable filaments as shown in Fig. S1b. (3) All instable filaments are additionally taken into account: The

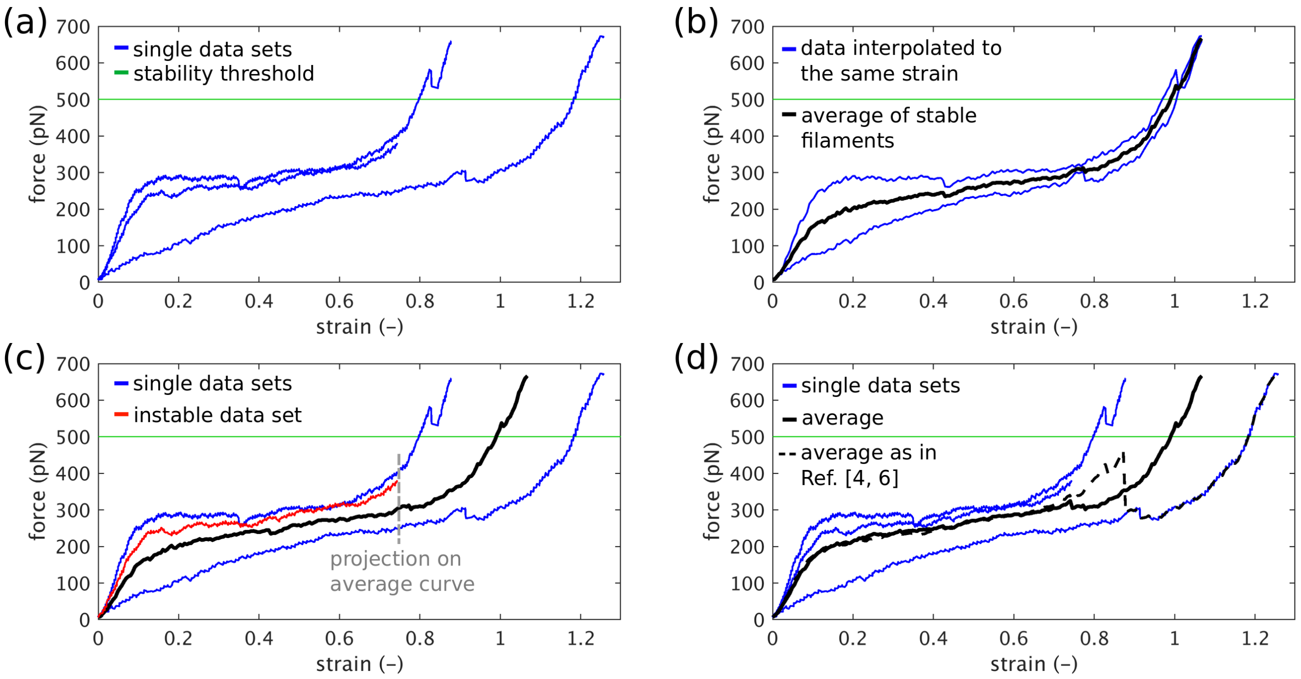


FIG. S1. (a-d) Force-strain averaging method for three typical force-strain data sets of single vimentin filaments in HB. (a) Typical force-strain data sets (blue) and stability threshold force (green). (b) Interpolation of stable force and strain data sets to 200 data points (blue) and average data (black). (c) Projection of the strain data of the instable filament on the average strain data calculated so far. (d) Average of all force data (black, continuous line) and average calculated with the method used in Refs. [4, 6] (black, dashed line).

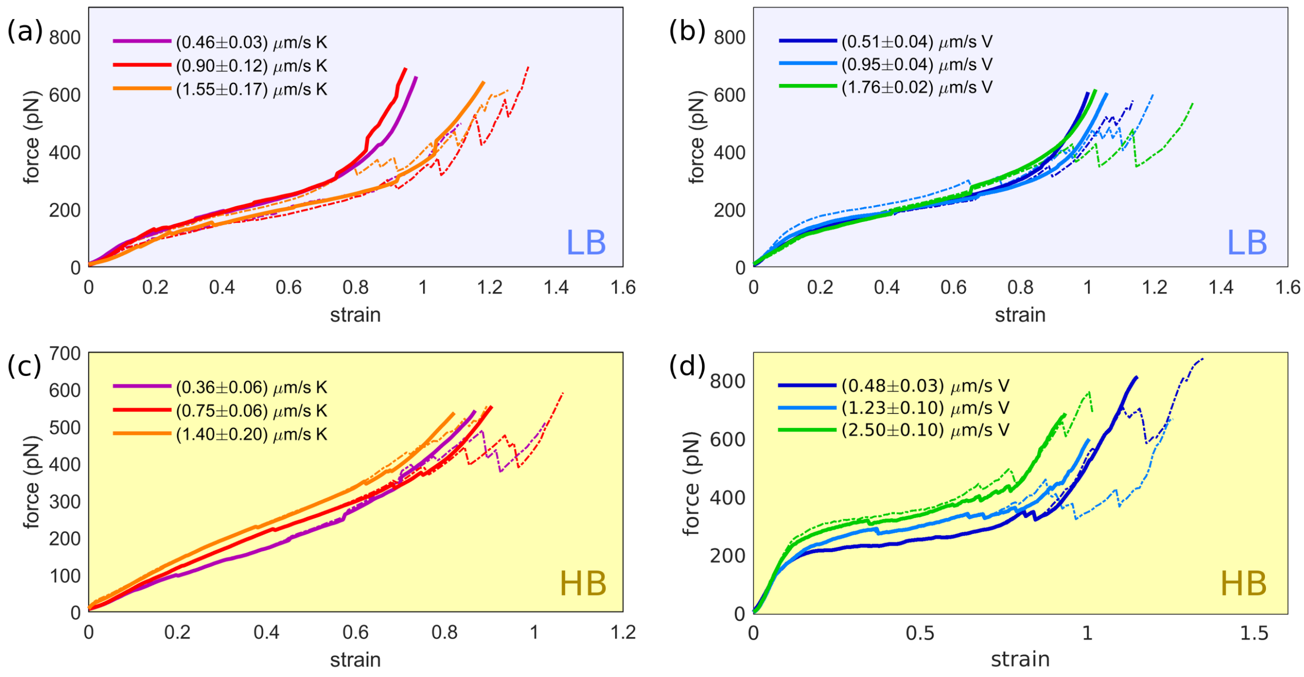


FIG. S2. (a)-(d) Force-strain curves comparing the force averaging method as in Refs. [4, 6] (dashed lines) and the the averaging method used here (continuous lines) for keratin and vimentin filaments in varying buffer conditions: (a) Keratin (K) filaments in LB, (b) vimentin filaments (V) in LB (blue background), (c) keratin filaments in HB, (d) vimentin filaments in HB (yellow background).

strain value of the average curve of all stable filaments closest to the maximum strain value of the instable filament is determined as shown in Fig. S1c. The force data of the instable filament are interpolated to the number of data points which are required to describe the average force data for stable filaments up to the point where the instable filament breaks. The force data of the instable filament are averaged with the average force curve for stable filaments for each strain value as in Fig. S1d. Since only one filament is instable, but the average force data are calculated from two stable filaments in steps (1) and (2), the data from the instable filament are weight as one third of the average of the two stable curves. The data from the instable filament are only taken into account up to the strain at which it breaks.

In Refs. [4, 6], the force data are averaged without calculating the average strain data. By contrast, the method presented here also takes the average strain into account, avoids kinks for high strain values and does not calculate the average force-strain curve from a few data curves only in the high strain regime. Both methods are compared in Fig. S1d and Fig. S2.

**Input energy calculation:** We integrate the force-strain curves within the force range of 0 to 500 pN using the MatLab function `trapz`. The results in units of $k_BT$ are shown in Fig. S3a.

**AFM height profile analysis:** The areas of interest in AFM height images are chosen manually. The selection is thresholded, so that the filament is clearly distinguishable from the background. Since the side length of the AFM images is merely 3-3.5 $\mu$m, single filaments do not form loops, so that it is possible to describe the filament contour as a function $y = L(x)$. The $x$-coordinate is obtained by averaging the positions of pixels which are occupied by the filament at $x$. The `smoothingspline`-fit function from MatLab is used to smooth the filament contour. The filament height is measured along perpendicular lines to the filament contour.

#### SIMULATIONS

Single keratin and vimentin IFs under tension are modeled with a mechanic model which is based on Refs. [4, 7, 8]. One unit length filament (ULF) has $N_P$ parallel monomers, which are arranged in parallel and all have the same length before loading. The monomers can be divided into $N_C$ $N_M$-mers. *E.g.*, a ULF with $N_P = 32$ monomers can be divided into 16 dimers as we assume in the simulation of vimentin filaments in LB ($N_C = 16$, $N_M = 2$) [9, 10]. In the following, we will assume two different coupling cases: In case 1, the $N_M$-mers act as independent units; in case 2, the $N_M$-mers are coupled to one another. Thus, in case 1, the subunits are allowed to slide past each other, but they are still laterally arranged. This also supports the idea of a protofilament structure of the filament after assembly in which the protofilaments can slide past each other [11–13]. The total number of ULFs connected in series in one filament before loading is $N_S$.

**Uncoupled case 1: Length extension of the ULF occurs when all monomers of a $N_M$-mer are in the $\beta$-state**. When the $N_M$-mers can elongate independently, the ULF elongates once all monomers in one $N_M$-mer are in the $\beta$-state. A sketch of the model is shown in the main text in Fig. 4a. In this case, all $N_M$-ers that are fully in the $\beta$-state are exposed to the loading force, so that they contribute to the ULF stiffness even if there are still $N_M$-mers in the $\alpha$-state in the same ULF.

The spring constant of an $\alpha$-helix is $\kappa_\alpha$, the spring constant of the monomer in the $\beta$-state $\kappa_\beta$ and the spring constant of all linkers is $\kappa_L$. $A_{j,m}$ is the number of monomers in the $\alpha$-state and $B_{j,m}$ is the number of monomers in the $\beta$-state in the $m$th $N_M$-mer in the $j$th ULF, thus $A_{j,m} + B_{j,m} = N_M$. $I_j$ is the number of $N_M$-mers of which all monomers are in the $\beta$-state. Thus, the spring constant becomes:

$$\kappa_j = \begin{cases} \left(\frac{1}{\kappa_L} + \frac{1}{N_P \kappa_\beta}\right)^{-1} & \text{for } \sum\limits_{m=1}^{N_C} A_{j,m} = 0 \\ \left(\frac{1}{\kappa_L} + \frac{1}{\sum\limits_{m=1}^{N_C} A_{j,m}\kappa_\alpha + N_M \kappa_\beta I_j}\right)^{-1} & \text{for } \sum\limits_{m=1}^{N_C} A_{j,m} > 0. \end{cases} \tag{1}$$

The spring constant of the entire filament is calculated by $\kappa_F = 1/(\sum_{j=1}^{N_S} 1/\kappa_j)$. Similar to Ref. [4], we neglect viscous and entropic contributions for simplicity.

The equilibrium reaction constant $K_{eq}$ for the $\alpha$ to $\beta$-state transition is:

$$K_{eq} = \frac{r^{\alpha\to\beta}}{r^{\beta\to\alpha}} = \exp(-\Delta G/(k_BT)) = 1/\gamma\,, \tag{2}$$

where we define $\gamma = \exp(\Delta G/(k_BT))$.

The load distribution factor $\theta$ ($0 < \theta < 1$) [14, 15] ensures detailed balance and energy conservation during opening and closing of a single $\alpha$-helix to the $\beta$-state and back. The force is distributed among the number of independent subunits $N_C$ and within these elements on the number of monomers in the $\alpha$-helix configuration $A_{j,m}$. The force $\phi = F/F_\alpha$ is dimensionless and scaled to the force $F_\alpha$ which is required to open an $\alpha$-helix into the $\beta$-state. The time $\tau$ is dimensionless and related to the time $t$ with the zero-force reaction rate from a monomer in the $\alpha$- to the $\beta$-state $r_0^{\alpha\to\beta}$ by $\tau = r_0^{\alpha\to\beta} t$. We assume Bell-Evans kinetics [7], so that the $\alpha$- to $\beta$-transition rate is:

$$r_{A_{j,m}}^{\alpha\to\beta} = A_{j,m} r_0^{\alpha\to\beta} \exp\left(\frac{\theta\phi}{N_C A_{j,m}}\right). \tag{3}$$

All monomers of one $N_M$-mer in the $\beta$-state fold back to the $\alpha$-state with the equilibrium constant since we assume that they are free of force if there are still $\alpha$-helices in the same $N_M$-mer, corresponding to the condition $A_{j,m} > 0$. If all monomers are in the $\beta$-state ($A_{j,m} = 0$), the reaction rate to the $\alpha$-state depends on the force. In this case, the force $(1-\theta)\phi$ is distributed to all $N_P$ monomers. Therefore, the closing of the $\beta$-states to the $\alpha$-helix becomes:

$$r^{\beta\to\alpha}_{A_{j,m}} = \begin{cases} r_0^{\alpha\to\beta} B_{j,m}\gamma & \text{for } A_{j,m} > 0 \\ N_M r_0^{\alpha\to\beta}\gamma \exp\left(\frac{-(1-\theta)\phi}{N_P}\right) & \text{for } A_{j,m} = 0. \end{cases} \tag{4}$$

From these rates, we calculate the probability $P_{A_{j,m}}$ that a certain number of a monomers is in the $\alpha$-helix configuration. This is vital for performing the Gillespie algorithm in the simulation.

$$\begin{aligned} \frac{dP_{A_{j,m}}}{dt} &= r^{\alpha\to\beta}_{A_{j,m}+1} P_{A_{j,m}+1} + r^{\beta\to\alpha}_{A_{j,m}-1} P_{A_{j,m}-1} \\ &\quad -(r^{\alpha\to\beta}_{A_{j,m}} + r^{\beta\to\alpha}_{A_{j,m}}) P_{A_{j,m}} \end{aligned} \tag{5}$$

When all monomers in one $N_M$-mer are in the $\beta$-state, the ULF extends by $\Delta L$. To make the simulation run dimensionless, we normalize $\lambda = L_0/\Delta L$, where $L_0$ is a characteristic length of the filament, *e.g.* the original filament length. Thus, in this case, the extension of the $j$th ULF $\lambda_j$ is:

$$\lambda_j = \begin{cases} 0 & \text{if for all } m\text{: } A_{j,m} > 0 \\ 1 & \text{if for any } m\text{: } A_{j,m} = 0. \end{cases} \tag{6}$$

The total extension of the filament then is $\lambda_{tot} = \sum_{i=1}^{N_S} \lambda_j$. Since the optical traps pull on the filament with a constant velocity $v$, the end-to-end distance is $x(t) = vt$. The force on the filament becomes

$$\phi = \kappa_F(x - \lambda_{tot}) \,. \tag{7}$$

**Coupled case 2: Length extension of the ULF occurs when all monomers of the ULF are in the $\beta$-state**. The probability that a certain number of monomers is in the $\alpha$-helical configuration is still calculated as in Eq. 5. The force calculation is the same as in Eq. 7. However, in this case, the force that acts on a ULF is distributed over all monomers in the $\alpha$-state in the ULF. The length of the filament increases only once all $\alpha$-helices in one ULF are in the $\beta$-state. It is no longer sufficient that one $N_M$-mer is in the $\beta$-state, as in case 1 above. This is equivalent to the assumption that subunits do not slide past each other. A sketch of this model is shown in the main text in Fig. 4b.

$N_M$-mers in the $\beta$-state do not contribute to the overall spring constant, if there is any monomer in the $\alpha$-state in the same ULF. Thus, the spring constant of the $j$th ULF is calculated as:

$$\kappa_j = \begin{cases} \left(\frac{1}{\kappa_L} + \frac{1}{N_P \kappa_\beta}\right)^{-1} & \text{for } \sum\limits_{m=1}^{N_C} A_{j,m} = 0 \\ \left(\frac{1}{\kappa_L} + \frac{1}{\sum\limits_{m=1}^{N_C} A_{j,m}\kappa_\alpha}\right)^{-1} & \text{for } \sum\limits_{m=1}^{N_C} A_{j,m} > 0 \end{cases} \tag{8}$$

For the $m$th subunit in the $j$th ULF we obtain:

$$r^{\alpha\to\beta}_{N_{j,m}} = A_{j,m} r_0^{\alpha\to\beta} \exp\left(\frac{\theta\phi}{\sum\limits_{m=1}^{N_C} A_{j,m}}\right) , \tag{9}$$

since the force $\theta\phi$ is equally distributed among all monomers in the $\alpha$-state $\sum_{m=1}^{N_C} A_{j,m}$.

For the reaction from $\beta$- to $\alpha$-state we get:

$$r^{\beta\to\alpha}_{N_{j,m}} = \begin{cases} r_0^{\alpha\to\beta} B_{j,m}\gamma & \text{for } \sum\limits_{m=1}^{N_C} A_{j,m} > 0 \\ N_M r_0^{\alpha\to\beta}\gamma \exp\left(\frac{-(1-\theta)\phi}{N_P}\right) & \text{for } \sum\limits_{m=1}^{N_C} A_{j,m} = 0. \end{cases} \tag{10}$$

The crucial difference to case (1) is that the filament extends in length *only* when all monomers within a ULF are in the $\beta$-state. Comparing to Eq. 6, we get:

$$\lambda_j = \begin{cases} 0 & \text{if for any } m: \; A_{j,m} > 0 \\ 1 & \text{if } \sum\limits_{m=1}^{N_C} A_{j,m} = 0. \end{cases} \tag{11}$$

As shown in Fig. S4a, a plateau in the force-strain data forms in the coupled case, but not for the uncoupled case for the same reasonable parameters (see Table I). With these parameters, we can classify the linkers as "soft transducers" since $(\kappa_\alpha + \kappa_\beta) \lesssim \kappa_L$ [16]. To show that the plateau is caused by the high lateral coupling strength between the subunits and not by the different number of subunits in keratin filaments ($N_P = 16$) and vimentin filaments ($N_P = 32$), the uncoupled case (1) is simulated for 32 monomers instead of 16 and shown as the "control" data in Fig. S4b. The same set of parameters for either 32 and 16 monomers per cross-section in the uncoupled case does not lead to a plateau. This supports our hypothesis that the lateral coupling strength between the subunits determines the force-strain behavior and not the number of monomers per cross-section.

**Fitting and rescaling to SI units:** The initial fitting parameters of the simulations are guessed so that they resemble the data. The `fminsearch`-function of MatLab is used to find the minimum sum $f$ of the absolute differences between the single values of the experimental data and the simulated data. The parameters $\kappa_L$,

TABLE I. Simulation parameters in simulation units.

| condition | case[a] | $N_P$ | $\Delta G$ | $\kappa_\alpha$ | $\kappa_\beta$ | $\kappa_L$ | $\Delta L$ | plateau |
|---|---|---|---|---|---|---|---|---|
| K in LB | 1 | 16 | $2.0 \pm 0.6$ | $4.4 \pm 1.9$ | $22 \pm 9$ | $64 \pm 7$ | $0.6 \pm 0.06$ | no |
| V in LB | 1 | 32 | $1.60 \pm 0.15$ | $2.1 \pm 0.7$ | $16 \pm 6$ | $18.0 \pm 1.9$ | $1.16 \pm 0.07$ | no |
| K in HB | 1 | 16 | $3.1 \pm 0.4$ | $4.6 \pm 1.3$ | $20 \pm 7$ | $60 \pm 6$ | $0.50 \pm 0.05$ | no |
| V in HB | 2 | 32 | $0.41 \pm 0.09$ | $11 \pm 4$ | $7 \pm 3$ | $27 \pm 3$ | $1.43 \pm 0.08$ | yes |
| V control | 1 | 32 | 3.1 | 4.6 | 20 | 60 | 0.5 | no |

[a] Uncoupled case 1 or coupled case 2.

TABLE II. Comparison of amino acids in K8, K18 and vimentin in at the positions vital for compaction. The charge pattern in vimentin allows compaction (red: positively charged amino acids, blue: negatively charged amino acids, green: amino acid with a large residue that can "open" the linker region for compaction) (adapted from Ref. [17]).

| protein | Positions L1 | Sequence: L1-1B | position L12 | Sequence: linker L12 |
|---|---|---|---|---|
| K8 | 127-136 | QQQKTARSNM-DNMFES | 238-254 | QISDTSVVLSMDNSRSL |
| K18 | 116-125 | LEKKGPQVRD-WSHYFK | 227-243 | QIASSGLTVEVDAPKSQ |
| Vimentin | 139-147 | KGQGKSRLG-DLYEEE | 250-264 | QEQHVQIDVDVSKPD |

$\kappa_\alpha$, $\kappa_\beta$, $\Delta G$ and $\Delta L$ are varied by `fminsearch` to find the minimum of $f$. To obtain the strain $\varepsilon$ in the units of the experiment, the strain $x$ of the filament in the simulation is rescaled with the relative length change $L_M/\Delta L$ of a monomer when all $\alpha$-helices open to the $\beta$-state, where the monomer length $L_M$ and $\Delta L$ are estimated from structural data [6] and $n$ is the average number of ULFs in the experiment represented by one ULF in the simulation. For the case of keratin filaments in LB this leads us to:

$$\varepsilon = \frac{x}{N_S L_M n / \Delta L} \approx 0.0059x \,. \qquad (12)$$

For the scaling factor of the force, we relate the change of the force $\Delta F$ to the length change as in Eq. 7, thus for keratin filaments in LB:

$$\Delta\phi = \kappa_F \Delta L \approx 11.8 pN \,, \qquad (13)$$

where $\kappa_F = 0.22$ pN/nm [6]. Thus, 1 simulation force unit corresponds to about 11.8 pN in the experiment. With the same rescaling procedure, we calculate the free energy difference $\Delta G$ between the $\alpha$- and $\beta$-state in SI units as presented in the main text.

The results for $\Delta G$ obtained from the fit agree well with literature: For example, 0.78 $k_BT$/amino acid in idealized model polyalanine $\beta$-sheets [18], 0.95 $k_BT$/amino acid in human amylin [19] and 1.26 $k_BT$/amino acid in a $\beta$-sheet formed by two alanine dipeptide molecules [20] were found.

In addition to the $\alpha$-helix–to–$\beta$-state transition free energy ($\Delta G$), we also estimate the spring constant of the

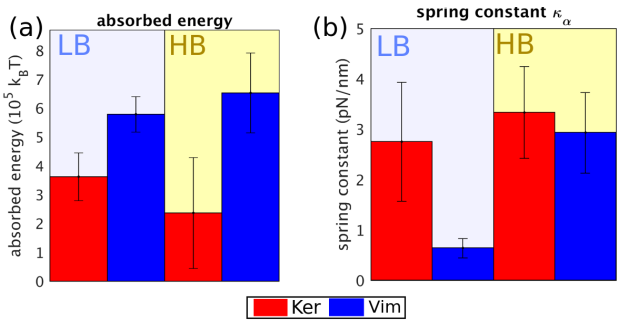


FIG. S3. (a) Input energy of keratin and vimentin filaments in two buffer conditions (blue background: LB, yellow background: HB). (b) $\alpha$-helical spring constants for keratin and vimentin filaments in two buffer conditions (LB, HB) determined from simulation parameters.

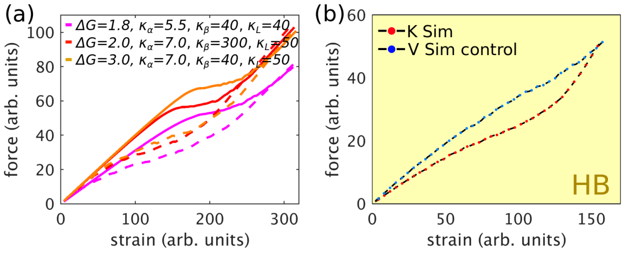


FIG. S4. (a) Simulation results for the same parameters for the uncoupled model (dashed lines) and the coupled model (continuous lines) for $\Delta L = 1$. (b) Comparison of the simulation of 32 parallel monomers (vimentin) with uncoupled subunits (blue-black, "control") and 16 parallel monomers (keratin) with uncoupled subunits (red-black). A plateau does not evolve merely as a consequence of an increased number of monomers.

$\alpha$-helices for keratin and vimentin filaments in the two different buffers as shown in Fig. S3b. Similar to the experimental data, we observe a pronounced stiffening of vimentin filaments in HB compared to LB. The $\alpha$-helical keratin stiffness increases slightly from LB to HB. An increased $\alpha$-helical stiffness in HB may be caused by additional intra-$\alpha$-helical attractions due to charge screening by the increased ion concentration.

#### FURTHER COMMENTS

**Compaction of vimentin IFs:** The phenomenon of compaction denotes the diameter reduction of IFs as a last step in assembly [2, 21]: The diameter of vimentin filaments shrinks from 17 nm to 9.5 nm [2, 22], whereas the diameter of keratin filaments changes from 10.9 nm to 9.4 nm [23], although vimentin IFs have twice as many monomers per cross-section on average. It seems to be vital that the transition of linker 1 (L1) to coil 1B contains amino acids, which are oppositely charged to the amino acids in linker 12 (L12), and that L12 contains a proline residue, as shown in Table II [17]. The oppositely charged amino acids attract each other and increase the coupling of neighboring subunits. The comparatively large proline residue allows for a spatially open arrangement of the linkers for the compaction step. By contrast, K8 does not contain a proline in linker L12 and similar charge patterns as in vimentin are not observed as shown in Table II. These molecular details are likely an explanation for compaction to occur in vimentin filaments, but not in K8/K18 filaments. This lack of compaction step in turn may be responsible for the weaker coupling between keratin subunits.

**Stability of $\alpha$-helices in keratin and vimentin:** As mentioned in the main text, the $\alpha$-helices in K8, K18 and vimentin have a highly similar stability. However, the number of polar or charged amino acids in coil 1A at positions, where hydrophobic amino acids are expected due to the $\alpha$-helical structure, is three times higher in keratin than in vimentin [24]; ions can stabilize this coil which may lead to increased stiffness. Yet, coil 1A is very short compared to the other coils (about 40 amino acids in coil 1A, 100 in coil 1B, 140 in coil 2) [24], so that the overall influence on stability from coil 1A may be comparatively small.

*

[14] A. Kolomeisky, Motor Proteins and Molecular Motors (CRC press, 2015).
